# Microbiota Modulation Induces Elevated Duodenal Eosinophils Upon Gluten Exposure in Mice: Implications for Non-Coeliac Gluten Sensitivity

**DOI:** 10.1101/2025.04.22.649964

**Authors:** Jennifer C. Pryor, Emily C. Hoedt, Wai Sinn Soh, Sophie Fowler, Shandelle Caban, Kyra Minahan, Simonne Sherwin, Cheenie Nieva, Huw McCarthy, Jay Horvat, Kateleen E. Hedley, Kerith Duncanson, Nicholas J. Talley, Grace L. Burns, Simon Keely

## Abstract

A growing proportion of the non-celiac population experience adverse symptoms to gluten. The pathogenesis of non-coeliac gluten sensitivity (NCGS) is unclear, but elevated duodenal eosinophils and altered mucosa-associated microbiota (MAM) populations have been reported. Given the microbiome’s role in gluten digestion and its susceptibility to antibiotics, we hypothesised that altering the microbiome with antibiotics would modify immune responses to gluten in mice. BALB/C mice consuming gluten-free chow received amoxicillin/clavulanate (5mg/kg) or PBS-vehicle daily for 5 days. Mice were then treated with a 3mg wheat-gluten suspension, or vehicle, on days 4 and 5 before sacrifice on day 7. Duodenal immune cells were analysed by histology and flow cytometry, while the duodenal MAM and faecal microbiome were characterised via 16S rRNA and shotgun metagenomic sequencing, respectively. Antibiotic treatment followed by gluten reintroduction significantly reduced *Staphylococcus* in the duodenal MAM, enriched *Bacteroides* in faeces, and resulted in altered microbial carbohydrate and lipid metabolism, compared to vehicle controls. Treatment with antibiotics and gluten also increased duodenal eosinophils which positively correlated with the genus *Blautia.* Flow cytometry revealed that antibiotics and gluten treatment resulted in a greater proportion of active eosinophils and epithelial γδ T-cells, compared to vehicle control mice. This study demonstrated that modulating the microbiome with antibiotics was sufficient to alter the immune response to gluten in mice. These findings suggest that the microbiome may determine the capacity for gluten to induce an immune response and offers a valuable insight into potential mechanisms underlying NCGS.

**New & Noteworthy:** A mouse model examined how microbial modulation affects immune responses to gluten. Antibiotic treatment followed by gluten reintroduction reduced duodenal *Staphylococcus* and altered microbial carbohydrate and lipid metabolism pathways in the faecal microbiome. Antibiotics and gluten treatment resulted in increased abundance and activation of duodenal eosinophils, and elevated γδ T-cells in the duodenal epithelium. These findings highlight the role the microbiome plays in gluten-induced immune responses, providing insights into mechanisms behind non-coeliac gluten sensitivity.

## Introduction

Wheat is a key food source for approximately 40% of the population, and gluten is its primary protein constituent [1]. Avoidance of wheat and adherence to a gluten free diet is increasing in the non-coeliac population; self-reports of wheat sensitivity approach 15% in Australia [2]. True non-coeliac wheat/gluten sensitivity (NCGS) is diagnosed following the exclusion of coeliac disease and immunoglobulin E (IgE)-mediated wheat allergy and is confirmed through a double-blind placebo-controlled wheat challenge [3]. People with NCGS report a range of symptoms associated with gluten consumption, including gastrointestinal (GI) symptoms such as nausea and bloating, as well as extra-intestinal symptoms like fatigue and brain-fog [4]. Despite the distinct symptom profile, no overt pathology is identified through blood tests or endoscopy in NCGS patients.

NCGS commonly coincides with disorders of gut brain interaction (DGBI), especially functional dyspepsia (FD), and irritable bowel syndrome (IBS) [4]. Whilst the aetiology remains poorly characterised, recent evidence describes an altered immune profile in NCGS patients. Within the intestinal epithelium, there is increased recruitment of lymphocytes [5, 6] and evidence of impaired barrier function [6, 7]. Furthermore, NCGS patients have increased duodenal eosinophils when compared to non-wheat sensitive controls [8], and mast cells have been found in higher proportions near mucosal nerve fibres [9]. As these cells are associated with allergic responses, they may contribute to symptoms in NCGS. Additionally, proinflammatory type one innate lymphoid cells (ILCs), are reportedly increased in the mucosa of NCGS patients [10]. Whilst these data indicate that an immune response has been generated in response to gluten, the mechanism that triggers this response remains unknown.

The primary protein and carbohydrate constituents of wheat are largely resistant to host derived enzymes and require the GI microbiome for digestion [11]. This process is key not only for the liberation of nutrients but may prevent inappropriate immune responses to incompletely digested foods [12]. Importantly, the GI microbiome in NCGS is altered compared to controls, with reports of increased duodenal *Actinobacillus* and elevated Ruminococcaceae in stool [13]. Moreover, *Blautia* is reduced in the stool of NCGS [14], aligning with similar findings in coeliac disease [15, 16]. An altered microbiome composition may affect digestion of wheat and intestinal barrier function which could enhance the capacity for gluten peptides to induce an immune response in NCGS patients [12, 17]. Interestingly, NCGS patients have increased serum biomarkers indicative of immune activation and intestinal epithelial damage. These biomarkers include soluble CD14 and lipopolysaccharide (LPS)-binding protein, which suggest an immune response to microbial peptides [7]. Additionally, increased levels of fatty acid-binding protein 2 (FABP2) indicate disruption of the intestinal epithelium [7]. Importantly, levels of these markers were observed to return to the level of controls following exclusion of gluten from the diet [7]. Antibiotics, used for the treatment of bacterial infections, non-specifically target both pathogenic and commensal microbes and thus altering the GI microbiome composition and function [18]. Multiple studies have implicated antibiotic treatment as a risk factor for the development of food allergies [19] and coeliac disease [20]; both conditions which involve an aberrant immune responses towards foods. Previous studies have shown that antibiotic treatment increases immune sensitisation towards food antigens in adjuvant-based mouse models of peanut allergy [21]. Thus, we hypothesise that modulating the GI microbiome with antibiotics will induce an aberrant immune response towards gluten peptides upon their introduction and we aimed to investigate this in a murine model.

## Methods

### Animal Model

#### Animal ethics and sourcing

All experiments were performed in accordance with the National Health and Medical Research Council Australian Code of Practice and approved by The University of Newcastle Animal Care and Ethics Committee (approval number A-2021-126). Female, BALB/c mice, were sourced from Australian BioResources Pty Ltd (Mona Vale, New South Wales (NSW), Australia) and were housed in individually ventilated cages, under specific pathogen free conditions. Mice were provided sterile, gluten-free chow (Cat. SF08-025; Specialty Feeds, Glen Forrest, Western Australia, Australia) and water *ad libitum,* commencing at the start of the 14-day acclimatisation phase and continuing for the duration of the study. All animals were maintained in a temperature (22°C ± 2°C) and humidity-controlled environment on a 12-hour light-dark cycle.

#### Murine model

Six-week-old BALB/c mice received daily 200µL oral gavage of amoxicillin/clavulanic acid (ABX; 5mg/kg; Sigma-Aldrich, St. Louis, Missouri, USA) or phosphate buffered saline (PBS) vehicle from days 0-4 (**Fig. 1A**). On days 4 and 5, mice were administered 3mg of gluten (Cat. G5004; Sigma-Aldrich), suspended in 0.02M acetic acid, or acetic acid vehicle alone, by oral gavage (200µL). Mice were briefly anaesthetised with 2% isoflurane (flow rate of 2L/min) prior to gavage with an 18-gauge ball end needle, attached to a 1mL syringe. On day seven, mice were sacrificed with an intraperitoneal injection of sodium pentobarbital (150mg/kg; Virbac, Carros, France).

**Figure 1:**
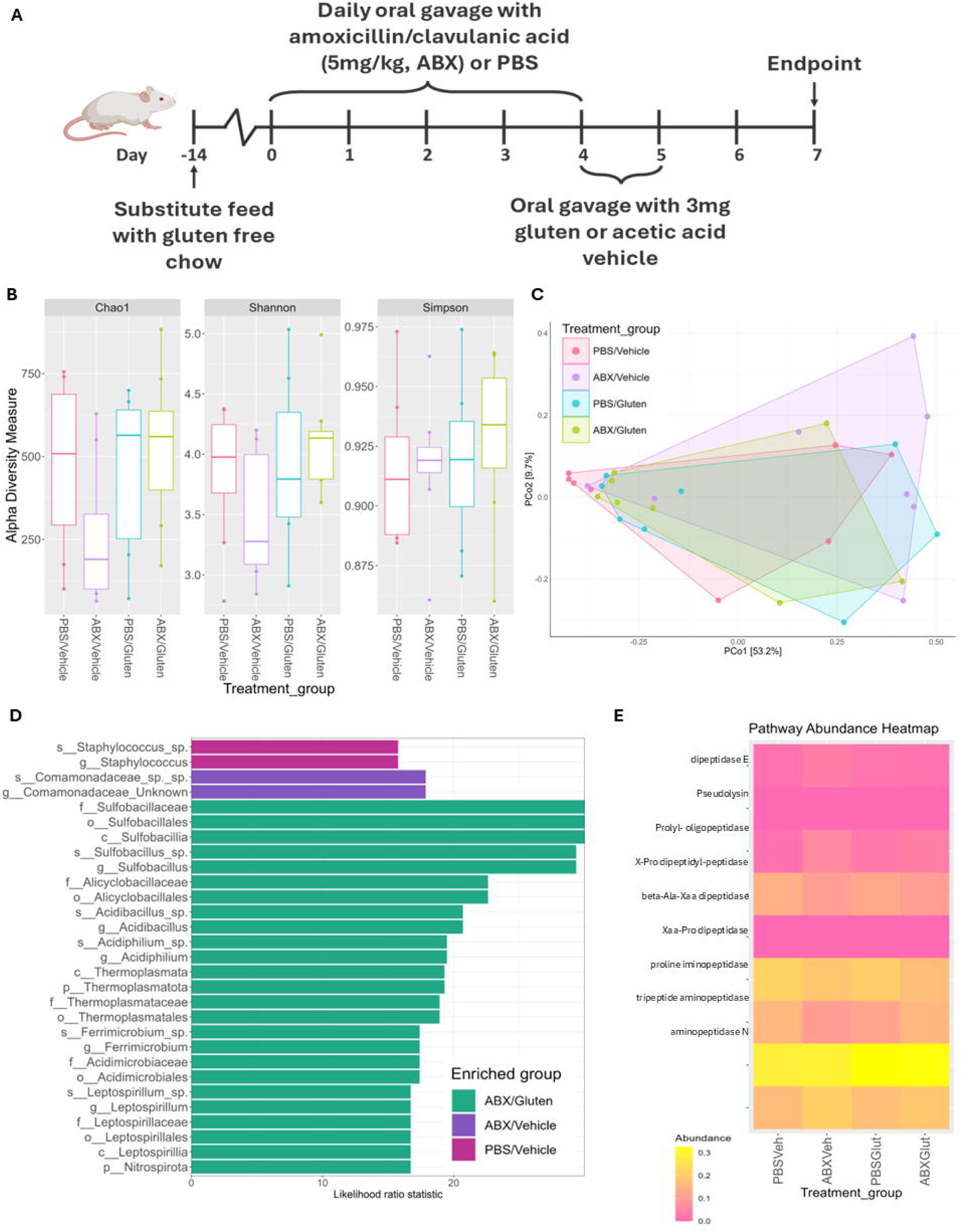
The duodenal mucosa associated microbiota of mice was assessed by 16S rRNA amplicon gene sequencing. **A)** Mice were treated with antibiotics (ABX) or phosphate buffered saline (PBS) for five days before oral gavage with gluten suspended in acetic acid vehicle or vehicle alone. **B)** Alpha diversity was measured by Chao1, Shannon and Simpson metrics. **C)** Beta diversity has been visualised with a PCoA plot of Bray-Curtis dissimilarity. **D)** Differentially abundant taxa were identified with EdgeR. **E)** Heatmap of predicted abundance of functional genes associated with gluten digestion. n=8/group (*p<0.05).

Based on preliminary data, an A priori power analysis using G*Power (v3.1.9.7) determined that a sample size of 5 animals per group was required to detect a significant difference between two independent means (Cohen’s d = 2.85, α = 0.05, power = 0.95). To account for control groups and experimental variability a final group size of n=8 for each analysis was chosen. There were four treatment groups, PBS/Vehicle, ABX/Vehicle, PBS/Gluten, and ABX/Gluten, and the experiment was repeated to ensure sufficient tissue for each analysis. At endpoint, tissue from the duodenal/jejunal junction and faeces collected from the colon were snap frozen and stored at –80°C for microbiome analysis (n=8/group). Duodenal tissue was also collected for histological analysis (n=8/group) or flow cytometric assessment (n=8/group).

### Microbiome Analysis

#### DNA Extraction from stool and small intestinal tissue

Genomic DNA was extracted from matched duodenal tissue and stool samples (n=8/group) using the Maxwell RSC Faecal Microbiome DNA Kit (Cat. AS1700; Promega, Wisconsin, USA), according to manufacturer’s instructions. A bead beating step was added during cell lysis using a Precellys homogeniser (Bertin Technologies, Montigny-le-Bretonneux, France) at 6000rpm for three, 60 second intervals. DNA was extracted in the Maxwell automated DNA purification instrument (Promega). Double stranded DNA was quantified with the Quantus Fluorometer using the QuantiFluor ONE dsDNA System (Cat. E4871; Promega). The presence of microbial DNA was confirmed by polymerase chain reaction (PCR) using primers to amplify the region between positions 1114 and 1221 of the 16S rRNA gene [22]. PCR products were separated on a 1% agarose gel containing 0.01% SYBR Safe DNA Gel Stain (Cat. S33102; Invitrogen) and visualised to confirm the presence of amplifiable microbial DNA.

#### 16S rRNA Sequencing of duodenal DNA

16S rRNA amplicon gene sequencing was performed on duodenal DNA to minimise the effects of host DNA contamination. Genomic DNA was submitted to the Australian Centre for Ecogenomics (ACE, Brisbane, Australia), for sequencing on the Illumina MiSeq system. Primers were selected to amplify the V3-V4 region (f-GAGTTTGATCNTGGCTCAG, r-GACTACHVGGGTWTCTAATCC).

Sequence data was processed, as previously described [23, 24], with the Quantitative Insights into Microbial Ecology 2 pipeline (QIIME2, v2022.2), where primers were removed with the Cutadapt plugin. The divisive amplicon denoising amplicon 2 (DADA2) package was utilised for quality assessment and denoising. Taxonomy of sequences was assigned with the SILVA reference database (v138) and the DADA2 package (v1.32.0) in RStudio (v2024.09.0) with R version 4.4.0, after which eukaryotic sequences and samples with fewer than 2000 total reads across all taxa were excluded. The R-package phyloseq (v1.48.0) was employed to evaluate alpha diversity using Chao1, Shannon, and Simpson metrics. Statistical significance of alpha diversity changes was determined using pairwise Willcox tests with Benjamin Huxburg p-value adjustment. Beta diversity was assessed using the ampvis2 package (v2.8.9) with Bray-Curtis distance, and significance was tested via permutational multivariate analysis of variance (PERMANOVA; adonis2 in vegan package v2.6.4). Differentially abundant taxa were investigated with LEfSe and EdgeR, within the microbiomeMarker package (v1.10.0). Taxonomy was correlated with histological findings using Spearman correlations in the microeco package (v1.9.1). The functional potential of the duodenal microbiota was predicted with PICRUSt2 within QIIME2 (v2021.11) and analysed using phyloseq (v1.48.0). The predicted relative abundance of KEGG orthologs, identified from literature as being related to the breakdown of gluten were compared between groups with Wilcoxon signed-rank test.

#### Shotgun metagenomic sequencing of faecal DNA

As there was little host DNA contamination in faecal samples, the faecal DNA was provided to Microba Pty Ltd (Brisbane, Australia), for shotgun metagenomic sequencing. Ahead of sequencing, libraries were prepared using the Nextera DNA Flex Library Preparation Kit (20018705) and indexed with IDT, as described previously [23]. The libraries were combined equally into a sequencing pool which was quantified and sequenced using the NovaSeq6000 platform with 2x 150bp paired-end chemistry, targeting a depth of 3Gb per sample, with a minimum depth of 2Gb.

Sequencing data was processed and analysed following previously published methods [23]. Briefly this consisted of trimming raw sequence data using the KneadData (v0.12.0), taxonomy assignment with MetaPhlAn4 (v4.0.6) [25], and functional assignments with HUMAnN3 (v3.8) [26], all using default settings. Processed data was then imported into RStudio for visualisation of alpha and beta diversity with packages phyloseq (v1.48.0), ampvis2 (v2.8.9), SIAMCAT (v2.7.2), microeco (v1.9.1), and microbiome (v1.26.0). Differentially abundant taxa were identified with EdgeR and LEfSe from the microbiomemarker (v1.10.0) package. Wilcoxon rank sum testing was performed on functional data, to identify differentially abundant pathways between each treatment group and the control, vehicle treated mice.

#### Histological Analysis of Immune Cells

Following euthanasia, the duodenum was collected (n=8 per group), and fat removed. The duodenum was cut open lengthways and prepared for histological assessment using the “Swiss Roll” technique [27]. Tissue was fixed in formalin for 48 hours and then transferred into 70% ethanol, prior to paraffin embedding by Core Histology Facility (Hunter Medical Research Institute, NSW, Australia). Tissue was sectioned at a thickness of 3µm and transferred onto slides for staining.

The tissue was deparaffinised with xylene and rehydrated with a series of incubations in ethanol dilutions prior to washing in distilled water. Slides were then stained with Harris haematoxylin (Cat. AHH-1L; ProSciTech, Kirwan, Queensland, Australia) and a 0.025% solution of Eosin Y (Cat. E4009; Sigma-Aldrich). Stained slides were visualised with a light microscope (Olympus CX33) at 40x magnification, by a blinded observer. Eosinophils were quantified across five high powered fields of view (HPF) and the average was calculated for each slide [28]. The average number of intraepithelial lymphocytes (IELs) and goblet cells per 50 enterocytes was calculated from five separate, randomly selected villi per slide [28, 29]. Additional slides were stained with Toluidine blue solution (1g/L, pH2.5; Cat. 52040; ProSciTech) for visualisation of mast cells. These slides were visualised at 20x magnification by a blinded observer and the average number of mast cells from five HPFs was calculated for each slide.

### Flow cytometry

#### Isolation of duodenal mucosal immune cells

At the experimental endpoint, duodenal tissue was collected and cut into 0.5cm pieces before being thoroughly washed in PBS. Duodenal mucosal immune cells were isolated as previously described, with minor modifications [28, 30, 31]. The epithelial digestion solution contained 1x Hanks’ balanced salt solution (HBSS; without Ca2^+^/Mg2^+^; Thermo Fisher, Waltham, Massachusetts, USA) with 10mM HEPES, 5mM EDTA, 1mM dithiothreitol, and 5% foetal calf serum (FCS). Tissue samples were incubated in the epithelial digestion solution for 15 minutes at a time and the flow throughs, which contain the IEL fraction, were combined and kept on ice. To isolate lamina propria lymphocytes (LPL), the tissue pieces were next incubated for 20 minutes in a lamina propria digestion solution of HBSS (containing Ca2+/Mg2+; Thermo Fisher) with 10mM HEPES, 0.5mg/mL collagenase D (Cat. 11088866001; Sigma-Aldrich), 0.5mg/mL DNase I (Cat. 10104159001; Sigma-Aldrich), 3mg/mL dispase II (D4693; Sigma-Aldrich), and 5% FCS [28, 31]. The IEL and LPL fractions were centrifuged at 350xg for 10 minutes and washed once with PBS. Finally, samples were resuspended in flow cytometry buffer (PBS, 2mM EDTA, 1% FCS) for staining.

#### Staining of duodenal immune cells for flow cytometry

The total LPL fractions from each animal were divided into two equal portions for staining with a panel of antibodies to detect either eosinophils or lymphocytes and ILCs. The total IEL fractions were stained with the lymphocyte and ILC antibody panel only. Cell suspensions were centrifuged and resuspended in fixable viability dye at a concentration of 1:1000. Following a 15-minute incubation, Fc Block (Cat. 564220; BD Biosciences, Franklin Lakes, New Jersey, USA) was added to a concentration of 1:1000 before a further 10-minute incubation. To identify eosinophils, LPL cell fractions were stained for 30 minutes at 4°C with antibodies to detect CD11b (BV421, BioLegend, San Diego, California, USA), CD274 (BV711, BioLegend), CD80 (FITC, BioLegend), CD62L (PE, BioLegend), Siglec-F (APC, BioLegend) and CD45 (BUV395, BD Biosciences). To characterise ILCs and lymphocytes, IEL and LPL cell suspensions were stained with antibodies towards the surface markers CD90.2 (APC-Cy7, BD Biosciences), CD45 (BUV395, BD Biosciences), CD127 (BV711, BD Biosciences), CD3 (PE-Cy7, BioLegend), γδ T-cell receptor (γδ-TCR; FITC, Biolegend) and lineage cocktail (APC, BD Biosciences), for 30 minutes at 4°C. Following staining, all cells were centrifuged, washed with PBS and resuspended in a fixation buffer (BioLegend), containing 4% paraformaldehyde, for 15 minutes. Once fixed, cells stained with the eosinophil antibody panel were washed with PBS, resuspended in flow cytometry buffer and stored at 4°C prior to acquisition.

Following fixation, cells stained with the ILC/lymphocyte antibody panel were washed with PBS and resuspended in 1x intracellular staining permeabilization buffer (Cat. 421002; BioLegend). Cells were centrifuged and washed a second time in permeabilization buffer. Cells were then stained with antibodies towards the intracellular markers GATA3 (PE-CF594, BD Biosciences), TBET (BV421, BD Biosciences) and RORγt (PE, BD Biosciences), for 20 minutes at room temperature. Cells were washed twice in permeabilization buffer and resuspended in flow cytometry buffer for storage at 4°C.

#### Flow cytometric analysis

Flow cytometry acquisition was performed on the LSRFortessa™ X-20 Cell Analyzer with FACSDiva software (BD Biosciences). Flow cytometry data was analysed with FlowJo software (v10.9.0). From the total live, single cell population, eosinophils were identified as CD45^+^ CD11b^+^ Siglef-F^+^. Resident eosinophils were identified as CD62L^+^, whereas inflammatory eosinophils were CD62L^-^. Active eosinophils were recognised as the double positive population for CD80 and CD274, whilst basal eosinophils were double negative. Also, from the population of live, single cells, lymphocytes were identified as CD45^+^CD3^+^, and the presence of the γδTCR on these cells was assessed. The total population of ILCs were detected as lineage^-^CD90.2^+^CD45^+^CD127^+^ cells. ILC subsets were further differentiated by the presence of transcription factors; TBET^+^ ILC1s, GATA3^+^ ILC2s, RORγt^+^ ILC3s. A detailed gating strategy is provided in **supplementary figure 1.**

### Statistics

Unless specified otherwise, graphs and statistical analyses were generated using Graphpad Prism 10, where data were presented as mean ± standard error of the mean (SEM). A significance threshold of p<0.05 was applied and statistically significant outliers were excluded from each dataset using the Graphpad Grubbs’ outlier test. Normality was assessed with the D’Agostino & Pearson test, where p>0.05 indicated normal distribution. For normally distributed data, an ordinary one-way ANOVA was used, with Fisher’s LSD test used for post-hoc correction. For non-normally distributed data, a non-parametric ANOVA (Kruskal-Wallis test) with Dunn’s test, was employed.

## Results

### Microbes associated with acidic environments are enriched in the duodenum of mice treated with both antibiotics and gluten

The duodenum is the first site where gluten peptides encounter the mucosal immune system following gastric digestion, making it a critical region for initiating immune responses. Given that this process can be influenced by the mucosa-associated microbiota (MAM), the duodenal MAM was sequenced in this study. A blank, technical reagent control was confirmed to have zero sequence reads, and the analysed samples had a mean read count of 5,814,444, containing a total of 6,758 taxa. Alpha diversity was evaluated using the Chao1, Shannon, and Simpson indices but no significant changes were observed (**Fig. 1B**). Similarly, beta diversity, analysed by Bray-Curtis dissimilarity, was not affected by treatments (p=0.203; **Fig. 1C**), indicating that the overall microbial community structure remained relatively stable despite the interventions.

Differential abundance tests revealed mixed results with LEfSe analysis not identifying any significantly enriched species, whereas EdgeR analysis identified several enriched taxa (**Fig. 1D**). Specifically, there was an enrichment of *Staphylococcus* in the vehicle-only treated group. In mice treated with antibiotics, there was an enrichment of Comamonadaceae. Mice treated with both antibiotics and gluten had the greatest number of significantly enriched microbes including, *Sulfobacillus, Acidibacillus, Acidiphilium, Ferrimicrobium*, and *Leptospirillum.* These species thrive in acidic environments, and many are involved in iron and sulphur cycling, suggesting a shift in metabolic activity within the duodenal MAM. To investigate the functional capacity of the duodenal MAM to breakdown gluten, the predicted relative abundance of nine KEGG orthologs relating to gluten-digesting peptidases [32] were compared between groups (**Fig. 1E**). The relative abundance of prolyl oligopeptidase (K01322) was significantly higher in mice treated with antibiotics only compared to vehicle-treated mice (p=0.02). Additionally, proline iminopeptidase (K01259) was more abundant in mice treated with both antibiotics and gluten, compared to those treated only with antibiotics (p=0.02).

### Histological assessment reveals duodenal eosinophils are increased in mice treated with both antibiotics and gluten

Duodenal sections were stained with haematoxylin and eosin stain to visualise to the tissue architecture and immune cell presence (**Fig. 2A**). Mice treated with both antibiotics and gluten had a significant increase in eosinophils, a key effector cell in allergic diseases, when compared to vehicle treated mice (mean ± SD: 10.7 ± 2.8 vs. 7.8 ± 2.9, respectively; p=0.02; n=8/group; **Fig. 2B**). There were no significant changes observed between treatment groups in IEL (**Fig. 2C**), or goblet cell counts (**Fig. 2D**) within the duodenum. Tissue was also stained with Toluidine Blue stain to quantify mast cells, however no significant change in mast cells was observed between groups (**Fig. 2E**).

**Figure 2:**
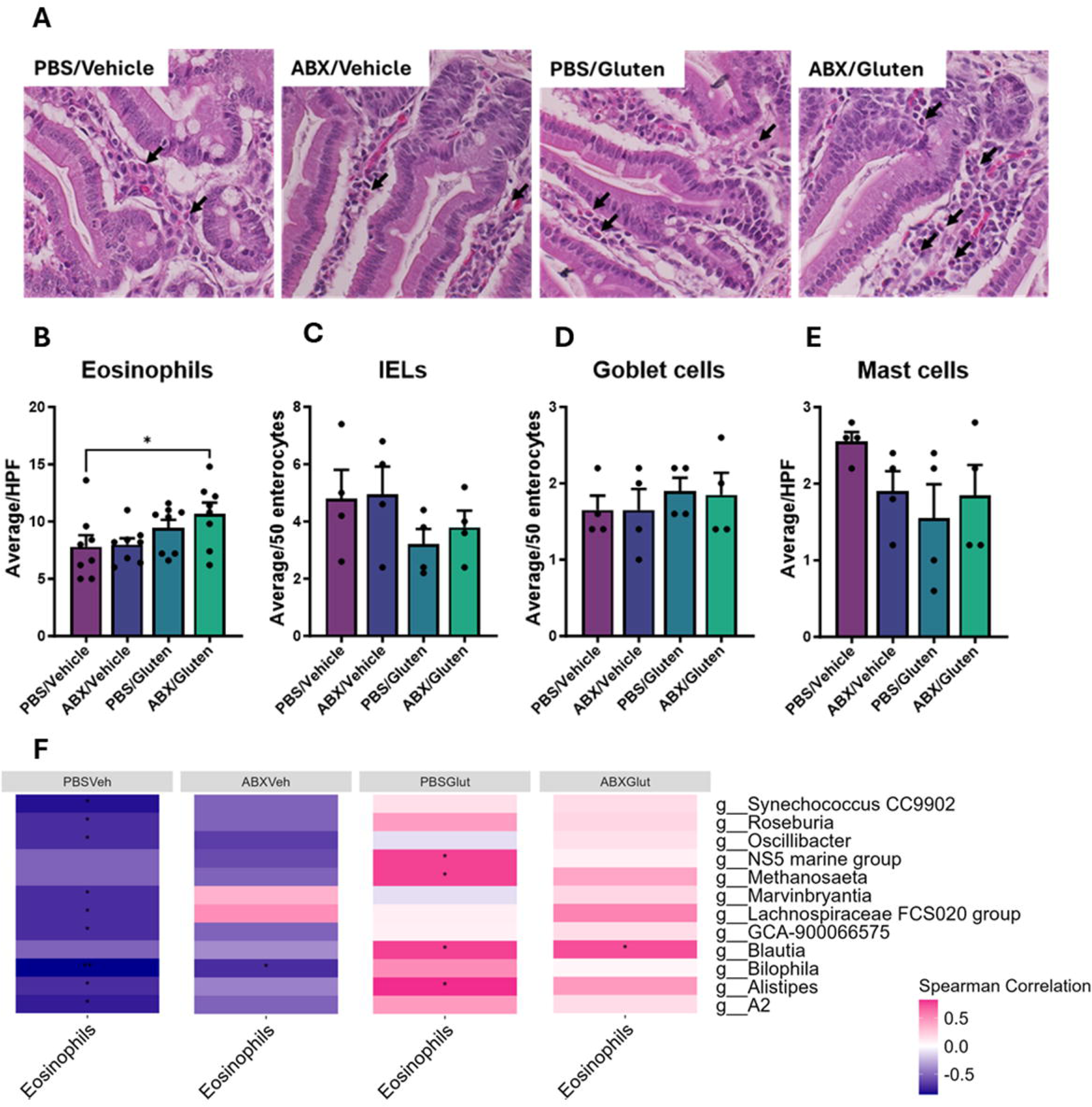
Histological assessment of mouse duodenums. Mice were treated with antibiotics (ABX) or phosphate buffered saline (PBS) for five days. Mice were then exposed to 3mg of gluten or vehicle by oral gavage. At experimental endpoint duodenal tissue sections were collected for staining and for microbiome analysis by 16S sequencing. **A)** Representative images of haematoxylin and eosin-stained tissue from each treatment group with eosinophils identified with arrows. **B)** The average number of eosinophils per high powered field (HPF) was quantified. **C)** The average number of intra-epithelial lymphocytes (IELs) per 50 enterocytes was assessed; **D)** as was the average number of goblet cells. **E)** Tissue was also stained with Toluidine Blue, and the average number of mast cells was calculated. **F)** A spearman correlation of duodenal microbes with histological eosinophils was conducted and is presented as a heatmap. Values are presented as mean ± SEM; n=4-8/group (*p<0.05).

Interestingly, correlations were observed between taxonomic 16S data and histological eosinophils in matched mouse samples (**Fig. 2F**). In vehicle treated control mice, lower duodenal eosinophil counts correlated with increased taxa belonging to genera *Roseburia, Oscillibacter, Marvinbryantia, Alistipes,* and *Bilophila. Bilophila* was also negatively correlated with eosinophil counts in mice treated only with antibiotics. Meanwhile, mice receiving gluten treatment alone, or in combination with antibiotics had a positive correlation between *Blautia* and duodenal eosinophils. Moreover, mice treated with only gluten had a positive correlation between eosinophils and the genera *Alistipes* and *Methanosaeta* (p<0.05).

### A greater proportion of lamina propria eosinophils are active following antibiotic treatment and reintroduction of gluten

Given the observed histological changes in eosinophils, we examined the proportion and phenotype of eosinophils in the duodenal, lamina propria by flow cytometry. This analysis did not identify changes to the proportion of live cells lamina propria cells (**Fig. 3A**), nor the total population of eosinophils (CD45^+^ CD11b^+^ Siglec-F^+^; **Fig. 3B**). Furthermore, there was no significant change in the proportion of inflammatory (CD62L^-^, **Fig. 3C**) or resident (CD62L^+^) eosinophils (**Fig. 3D**). There was however, a significant increase in the proportion of active eosinophils (CD80^+^CD274^+^ [33]) in mice treated with antibiotics and gluten compared to mice treated only with vehicle (mean ± SD: 3.0±2.7 vs. 1.4±0.6, respectively; p=0.04; n=8/group, **Fig. 3E**). Correspondingly, antibiotic and gluten treated mice had a significant reduction in basal (CD80^-^CD274^-^) eosinophils (80.9±13.6 vs. 90.2±3.9; p=0.02; n=8/group, **Fig. 3F**).

**Figure 3:**
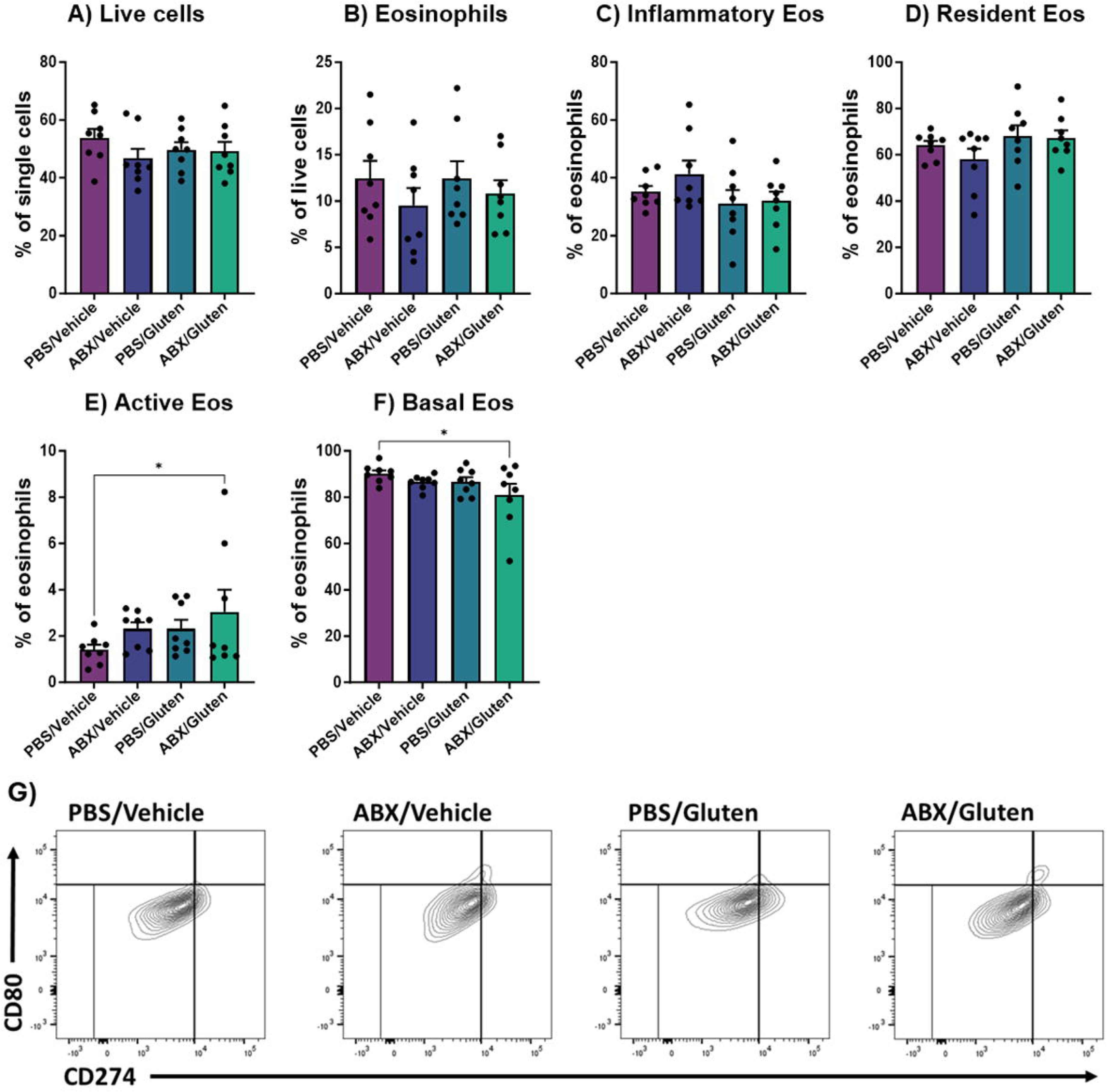
Mouse, duodenal, lamina propria eosinophils were assessed by flow cytometry. Mice were treated with antibiotics (ABX) or phosphate buffered saline (PBS) for five days before introduction of gluten or vehicle. At the experimental endpoint, duodenums were collected and enzymatically digested to liberate the lamina propria immune cells, which were then assessed my flow cytometry. **A)** The proportion of live cells in each group is shown. **B)** Total eosinophils (eos) were identified as CD45+ CD11b+ Siglec-f+ cells. The proportion of **C)** CD62L-inflammatory eosinophils, **D)** CD62L+ resident eosinophils, **E)** CD80+ PD-L1+ Active eosinophils, and **F)** CD80-PD-L1-Basal eosinophils are shown. **G)** Representative contour plots of eosinophils gated against CD80 and CD274 to identify active and basal eosinophils. Values are presented as mean ± SEM; n=8/group (*p<0.05).

### γδ T-cells and innate lymphoid cells are increased in the duodenal epithelium of antibiotic and gluten treated mice

The epithelial fraction of the duodenum was assessed by flow cytometry, and live cells were quantified. Mice receiving either antibiotics alone or gluten alone had a reduced proportion of live cells when compared to vehicle treated mice (p=0.03, p=0.0007, respectively; n=8/group; **Fig. 4A**). Despite these individual effects, there was no significant difference in viability of cells from mice receiving both antibiotics and gluten compared to vehicle treated mice. As epithelial lymphocytes are known to be altered in some gluten-related disorders, their abundance was investigated in this model. Lymphocytes, identified as CD45^+^CD3^+^ cells, were unchanged in the duodenal epithelium of mice (**Fig. 4B**). However, the proportion of γδ T-cells was significantly elevated in mice treated with both antibiotics and gluten when compared to vehicle control mice (8.6±2.5 vs. 5.8±1.7; p=0.05; n=8/group, **Fig. 4C**).

**Figure 4:**
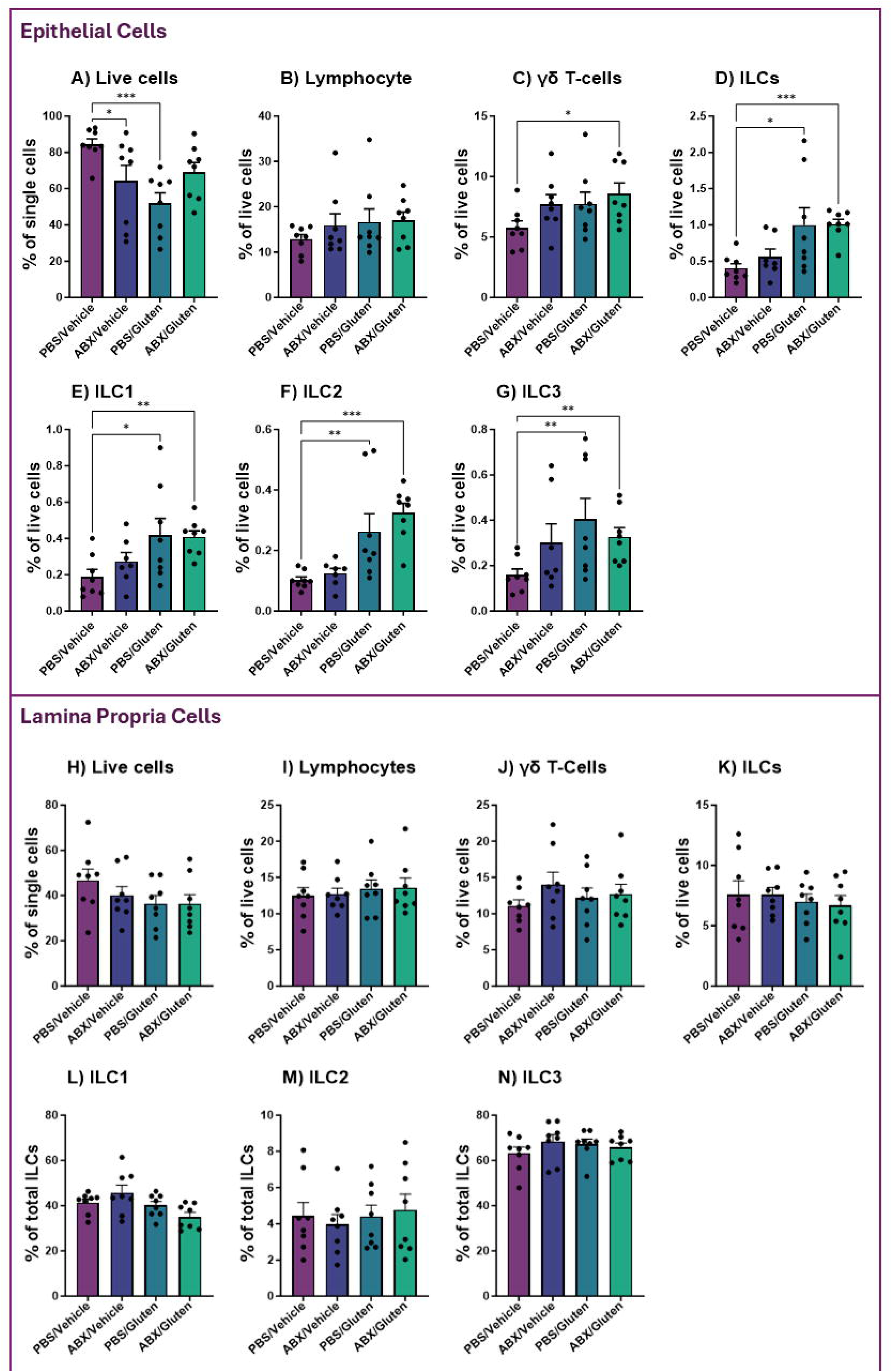
Duodenal lymphocytes and innate lymphoid cells (ILCs) were assessed in mice treated with antibiotics and gluten. Mouse duodenums were enzymatically digested to isolate the epithelial and lamina propria fractions and analysed by flow cytometry. The proportion of epithelial **A)** live cells, **B)** CD45^+^ CD3^+^ lymphocytes, and **C)** γδ TCR+ lymphocytes was assessed. Additionally, **D)** the proportion of total epithelial ILCs (lineage CD90.2^+^ CD45^+^ CD127^+^) was quantified. ILC subsets **E)** TBET^+^ ILC1 cells, **F)** GATA3^+^ ILC2 cells, and **G)** RORyt^+^ ILC3 cells were also analysed. The following lamina propria immune cell populations were also quantified: **H)** Live cells, **I)** CD45^+^ CD3^+^ lymphocytes, **J)** γδ TCR+ lymphocytes, **K)** total ILCs, **L)** ILC1s, **M)** ILC2s, and **N)** ILC3s. Values are presented as mean ± SEM; n=8/group (*p<0.05, **p<0.01, ***p<0.001).

ILCs are crucial in immune defence and maintaining intestinal homeostasis, thus their proportion was analysed within the duodenal epithelium. There was a significant increase in the proportion of ILCs (lineage^-^ CD90.2^+^ CD45^+^ CD127^+^) in the epithelium of mice treated with gluten only (1.0±0.7 vs. 0.4±0.2; p=0.01; n=8/group) or both antibiotics and gluten (1.0±0.2 vs. 0.4±0.2; p=0.0004; n=8/group), when compared to vehicle mice (**Fig. 4D**). This trend was similarly reflected in each of the assessed subsets of epithelial ILCs. When assessed as a proportion of live cells, there was a significant increase in TBET^+^ ILC1s (**Fig. 4E**) in gluten treated (0.4±0.3 vs. 0.2±0.1; p=0.02) and antibiotics & gluten treated mice (0.4±0.1 vs. 0.2±0.1; p=0.003), compared to vehicle treated mice. GATA3^+^ ILC2s (**Fig. 4F**) were also elevated in the epithelium of mice treated with gluten only (0.3±0.2 vs. 0.1±0.03; p=0.005), or antibiotics and gluten (0.3±0.09 vs. 0.1±0.03; p=0.0001). Similarly, RORγt^+^ ILC3s (**Fig. 4G**) were elevated in both gluten treated (0.4±0.3 vs. 0.2±0.07; p=0.009) and antibiotics and gluten treated mice (0.3±0.1 vs. 0.2±0.07; p=0.007), compared to control mice. Interestingly, none of the populations altered in the duodenal epithelium were disrupted in the lamina propria (**Fig. 4H-N**). The total cell counts for the epithelial and lamina propria cell populations analysed are provided in **supplementary figures 2 and 3** respectively.

### Species linked with digestion of complex carbohydrates are enriched in faeces of antibiotics and gluten treated mice

We next conducted shotgun metagenomic sequencing on faecal DNA to characterise the GI microbiome as a whole, and to understand the impact of antibiotics and gluten treatments. Samples sequenced at a depth of 3GB had a mean read count of 17,861,564 and a total of 1092 taxa. A blank reagent technical control was confirmed to have zero sequence reads. No significant differences in alpha (**Fig. 5A**) or beta diversity (p=0.081, n=8/group; **Fig. 5B**) were observed between treatment groups. Differential abundance analysis with EdgeR identified a range of significantly enriched taxa in each treatment group, most of which were unclassified (**Fig. 5C**). In the vehicle only treated mice, the enriched taxa largely belonged to the phylum Firmicutes and Bacteroidetes. Amongst the uncultured microbes, Lachnospiraceae bacterium 10-1 was enriched in mice treated with gluten only. Mice treated with only antibiotics exhibited an enrichment in the families Atopobiaceae and Streptococcaceae, and the genus *Lactococcus*. Treatment with both antibiotics and gluten resulted in an enrichment of *Bacteroides congonensis*, *Luxibacter massiliensis*, *Bacteroides uniformis*, and *Enterocloster aldenensis*. *Bacteroides* spp. are known for their ability to digest complex carbohydrates and produce short chain fatty acids (SCFAs). Interestingly, LEfSe analysis only identified significantly enriched microbes in the treatment group receiving gluten alone. The enriched taxa were from the family Muribaculaceae, the genus *Paramuribaculum,* and the species *Paramuribaculum intestinale*.

**Figure 5:**
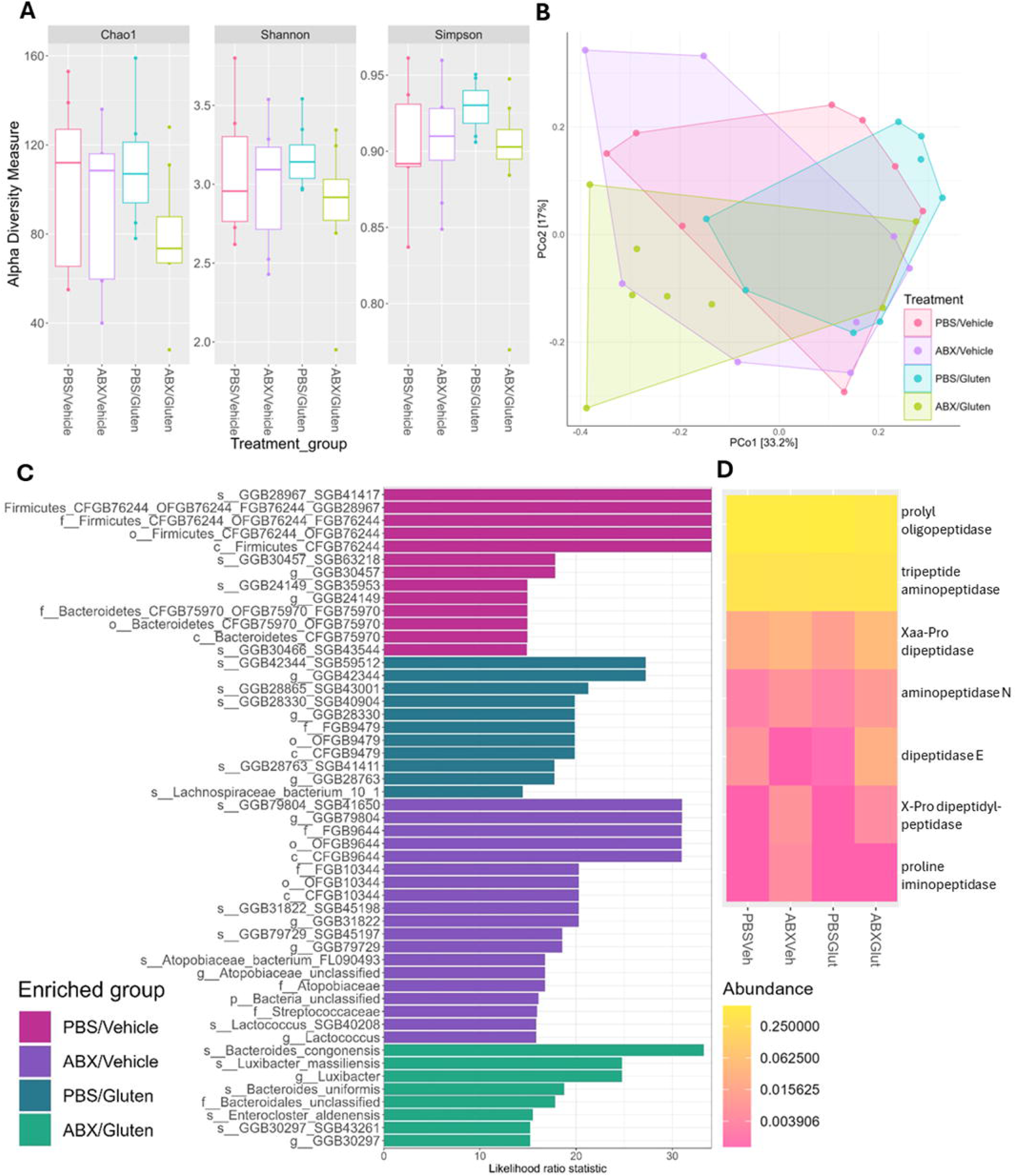
Shotgun metagenomic sequencing of mouse faecal microbiota. Mice received antibiotics (ABX) or phosphate buffered saline (PBS) before subsequent oral gavage with gluten or vehicle. Stool was collected at endpoint and the microbiome assessed by shotgun sequencing. **A)** Alpha diversity was assessed with Chao1, Shannon and Simpson. **B)** PCoA plot of Bray-Curtis dissimilarity for beta diversity visualisation. **C)** Differentially abundant taxa when measured by EdgeR. **D)** Heatmap of functional genes associated with gluten digestion. n=8/group.

Although gluten is typically digested in the small intestine, we examined the functional capacity of the faecal microbiome to breakdown gluten to understand the overall potential of the GI microbiome to digest gluten. KEGG orthologs relating to gluten-digesting peptidases were compared between treatment groups, and two of the orthologs present in the duodenal microbiome were absent from faecal samples. The relative abundance of prolyl oligopeptidase (K01322) was significantly reduced in mice treated with antibiotics and gluten when compared to mice treated only with gluten (p=0.003; **Fig. 5D**). No other significant changes in KEGG pathway abundance were observed compared to the antibiotics and gluten-treated mice.

### Pathways related to carbohydrate and lipid metabolism are altered in faeces of antibiotics and gluten treated mice

Wilcoxon rank sum testing was used to identify pathways in each of the treatment groups which differed significantly from vehicle treated control mice (**Fig. 6**). Across all treatment groups, a total of 57 pathways were significantly altered (p<0.05). In mice treated with antibiotics only, there was changes to pathways related to sulphur metabolism including an increase in L-methionine biosynthesis (PWY.5345), and reduced sulphate assimilation, reduction and cysteine biosynthesis related pathways (SULFATE.CYS.PWY; PWY1ZNC.1; PWY.821). There was also increased abundance of pathways related to energy production and the tricarboxylic acid cycle (PWY.5690; P105.PWY). Antibiotics only treated mice also had a reduction to pathways involved in biosynthesis of the amino acid L-phenylalanine (PWY.6628) and degradation of chitin (PWY.6906).

**Figure 6:**
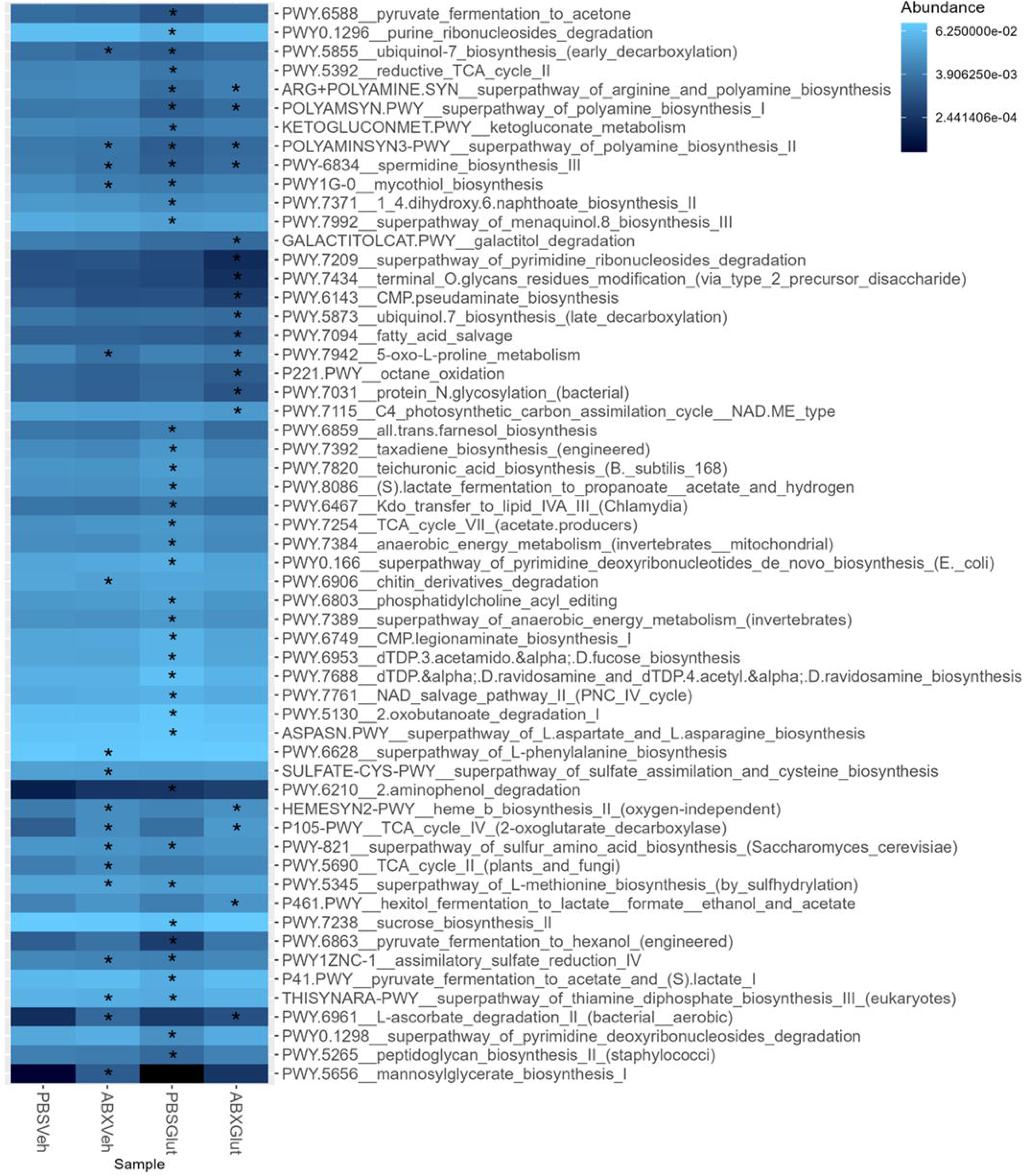
Significantly altered pathways in the faecal microbiome of antibiotic and gluten treated mice. Mice were treated daily for five days with antibiotics (ABX) or phosphate buffered saline (PBS), before being treated with 3mg of gluten (Glut) or the acetic acid vehicle (Veh) that the gluten was suspended in. Stool collected from the mice at endpoint was analysed by shotgun metagenomic sequencing. Pathways in each treatment group that significantly differed in abundance from PBS/Veh control mice were identified by Wilcoxon rank sum testing. Significantly altered pathways are shown in a heatmap with * indicating p<0.05. n=8/group.

Mice treated solely with gluten possessed some unique changes to faecal microbial pathways, including altered nucleotide metabolism, with an elevation in deoxyribonucleotide biosynthesis (PWY0.166) and reduction in deoxyribonucleosides and purine ribonucleosides degradation (PWY0.1298; PWY0.1296). There was also reduced pyruvate fermentation pathways (P41.PWY; PWY.6588; PWY.6863) and sucrose biosynthesis (PWY.7238). Further, these mice exhibited an increase in lactate fermentation (PWY.8086), anaerobic energy metabolism pathways (PWY.7384; PWY.7389), and nicotinamide adenine dinucleotide (NAD+) salvage pathway (PWY.7761).

Mice treated with gluten alone or in combination with antibiotics possessed a reduction to polyamine biosynthesis pathways (ARG.POLYAMINE.SYN; POLYAMSYN.PWY). Meanwhile antibiotics alone or in combination with gluten treatment resulted in enrichment to pathways of haem biosynthesis (HEMESYN2.PWY) and vitamin C degradation (PWY.6961) and a reduction in 5-oxo-L-proline metabolism (PWY.7942).

Finally, we observed some distinct pathways which were altered in mice treated with both antibiotics and gluten, such as a reduction in fatty acid salvage (PWY.7094). There is also altered carbohydrate metabolism with a reduction in galactitol degradation pathways (GALACTITOLCAT.PWY) and increased hexitol fermentation (P461.PWY). Antibiotics and gluten treated mice also exhibit a reduction in pathways for glycosylation (PWY.7031; PWY.7434) and nucleotide metabolism (PWY.7209). Thus, the combined treatment of antibiotics and gluten disrupts key microbial processes involved in energy production, carbohydrate utilisation, and macromolecule biosynthesis.

## Discussion

NCGS represents a significant health burden as it affects a substantial portion of the population, yet the underlying mechanisms remain poorly understood [34]. Current evidence suggests that individuals with NCGS experience alterations in both immune function [5–8] and GI microbiome composition [13, 14]. Importantly, one study observed increased markers suggestive of immune activation towards microbial components in NCGS, which were normalised by adherence to a gluten free diet [7]. This suggests an interplay between dietary gluten, the immune system, and the GI microbiome in the pathogenesis of NCGS.

While animal models present valuable opportunities to investigate the potential mechanisms driving NCGS development, to date murine models have largely focused on the pathogenesis of coeliac disease and wheat allergy [35]. When sensitised intraperitoneally with gliadin, transgenic mice expressing human leukocyte antigen (HLA) –DQ8 (the HLA allele associated with coeliac disease [36]) have heightened anti-gliadin antibody production, small intestinal IELs, and T-cell proliferation, reflecting the coeliac disease phenotype [37, 38]. Transfer of T-cells from gliadin sensitised mice to Rag1 knockout mice, which lack lymphocytes, results in duodenitis similar to that of coeliac disease patients, upon challenge with oral gluten [39]. Adjuvant based models of wheat allergy have shown that sensitisation to gliadin using aluminium hydroxide, with subsequent gliadin challenge can promote an IgE response and production of T helper (Th) type 2 cytokines in Balb/c mice [40]. A rat model exploring NCGS observed that repeated oral gavage with gliadin can result in mild inflammation including increased plasma interleukin (IL)-6, interferon gamma (IFNγ) and tumour necrosis factor α (TNFα); as well as elevated intestinal permeability as assessed by urinary lactulose/mannitol ratio [41]. However, existing models have largely failed to recapitulate mucosal changes and clinically relevant features of NCGS, including elevated duodenal eosinophils and altered IEL populations, without the use of adjuvants.

In this study, we demonstrated that modulation of the GI microbiome through antibiotic treatment was sufficient to alter the immune response to gluten in mice and promoted a duodenal immune phenotype that reflects aspects of NCGS. Our findings revealed several key immunological changes in mice treated with both antibiotics and gluten, including increased numbers of duodenal eosinophils, when assessed histologically, and a greater proportion of eosinophils expressing an activated phenotype. This aligns with work by Carroccio *et al.,* [8] where duodenal and rectal eosinophils were found to be elevated in NCGS patients when compared to controls, but not when compared to patients with coeliac disease. Additionally, we know that NCGS exists in a significant proportion of patients with FD [2], and duodenal eosinophils are increased in FD patients [42]. Eosinophils are effector cells of the immune system, which have been linked to a variety of conditions where foreign antigens trigger symptoms including eosinophilic oesophagitis, inflammatory bowel disease and asthma [43]. Activation of eosinophils can cause degranulation and release of pro-inflammatory factors such as major basic protein and eosinophil cationic protein [44–46], thus their elevation in NCGS and FD may prompt symptoms.

Patients with NCGS have been reported to have elevated epithelial lymphocytes [5, 6], however, in our mouse model we did not observe a change in the total epithelial CD3+ lymphocyte population. We did nevertheless observe increased populations of γδ T-cells in the duodenal epithelium of mice treated with both antibiotics and gluten. Epithelial γδ T-cells play an important role in immune surveillance and defence against pathogenic microbes [47]. γδ T-cells are significantly elevated in the duodenal epithelium of coeliac disease patients and have also been reported to be subtly increased in NCGS, when compared to controls [48]. Interestingly, an animal model of antibiotic treatment found that γδ T-cells were depleted in mice treated with antibiotics and this was associated with impaired oral tolerance to ovalbumin [49]. This suggests that our observed elevation in γδ T-cell numbers is gluten-dependant. Furthermore, gluten-treated mice in our study had a significant increase in epithelial ILCs. ILCs are often considered the innate counterparts to T-cells as they release cytokines that mimic some T-cell functions, thus they may be involved in recruiting effector cells in this model [50, 51]. Overall, the altered recruitment of immune cells to the duodenal epithelium of mice treated with antibiotics and gluten suggests an altered capacity to induce immune responses towards luminal antigens.

The relationship between the microbiome and immunological findings was important in this study. We observed that eosinophil numbers showed a positive correlation with duodenal *Blautia* in gluten-treated mice, while demonstrating a negative correlation with *Bilophila* in vehicle and antibiotic-treated mice. *Blautia* has previously been described to be increased in the duodenum of coeliac disease patients, but reduced in the faeces of coeliac [52] and NCGS patients [14]. Interestingly, a study of eosinophilic chronic rhinosinusitis has also described a positive correlation between mucosal *Blautia* and tissue eosinophil number [53]. The *Blautia* genus have largely been described as beneficial bacteria, due to the ability to produce short chain fatty acids (SCFA) and stimulate mucus growth [54], however this work suggests that the role of *Blautia* in eosinophilic disease requires further exploration. Although we observed no significant changes in alpha or beta diversity of either the duodenal or faecal microbiome, specific taxonomic changes were evident across treatment groups. We observed that *Staphylococcus,* a genus common in the GI microbiome, was reduced in the duodenal MAM of antibiotics and/or gluten treated mice compared to control mice. *Staphylococcus* is reduced in the duodenal microbiome of patients with coeliac disease and numerous species are described to have gluten degrading capacity, suggesting a possible role for *Staphylococcus* in determining the ability for gluten to promote immune responses [55, 56]. In the duodenal MAM, we also observed significant enrichment of genera *Sulfobacillus, Ferrimicrobium* and *Acidibacillus* in mice treated with both antibiotics and gluten. Whilst relatively little is known about these bacteria in respect to colonisation of the human gastrointestinal tract, their presence could suggest altered metabolic activity, given their roles in sulphur and iron oxidisation [57–59]. Relatedly, we also observed that the faeces of mice treated with antibiotics and gluten had an enrichment in haem biosynthesis. Given the critical role of iron in regulating immune and microbial homeostasis [60], and its known impact on the development of allergic diseases [61], its potential involvement in gluten sensitivity warrants further investigation.

Analysis of the faecal microbiome revealed that combined treatment with antibiotics and gluten led to enrichment of several specific species, including numerous *Bacteroides spp*. Given their carbohydrate degrading capacities, *Bacteroides* may contribute to the altered carbohydrate metabolism and glycosylation pathways observed in the functional faecal microbiome data. Whilst previous studies have shown that *Bacteroides uniformis* can improve inflammation in a colitis mouse models [62], there is also evidence that this species is elevated in the stool of coeliac disease patients [14]. Furthermore, we observed changes in faecal microbiome pathways related to SCFA production in mice treated with antibiotics and gluten. These included reduced fatty acid salvage and increased hexitol fermentation. SCFAs are released from microbial fermentation of indigestible food components and are important in maintaining GI health [63]. It has been reported that faecal SCFA content is higher in coeliac disease and patients with IBS compared to controls [64, 65]. Few studies have assessed SCFAs in NCGS, however, one has observed a reduction in faecal acetate levels in patients, while total SCFA concentrations remained unchanged compared to controls [66]. Future work employing metabolomics to assess SCFA content in this model would be a valuable addition. The distinct microbial signatures associated with antibiotic treatment and gluten exposure, suggest potential mechanistic links between microbiome composition and gluten sensitivity.

One limitation of this study is the reliance on 16S rRNA sequencing to characterise the duodenum MAM and predict the function capacity. Due to the high level of host DNA contamination in tissue samples, we were unable to perform shotgun metagenomic sequencing, which would have provided a more comprehensive analysis of the microbial community and its functional capabilities. To overcome this limitation, we conducted shotgun metagenomic sequencing on faecal samples instead, however future studies could utilise host DNA depletion, duodenal mucosal swabs [23], or culturomics [67] to allow for duodenal shotgun metagenomic sequencing.

The observed changes in the immune profile within the duodenal epithelium suggests future investigations into epithelial integrity in this model are required. Ussing chamber experiments of duodenal epithelial permeability and quantification of protein markers of barrier function would provide needed insights into the effect of antibiotics and subsequent gluten exposure on epithelial barrier function. Furthermore, the observed elevation in duodenal eosinophils warrants investigation into its underlying mechanisms. Future studies could examine Th cell subsets, particularly Th2 cells, and cytokine production to elucidate inflammatory pathways and drivers of eosinophil recruitment. Although we did not identify significant alterations in microbial pathways specifically associated with gluten-digesting capacity, we suggest that future work should utilise mechanistic assays to help clarify the role of microbial digestion of gluten in this model. The microbial enzymatic activity of mouse duodenal washes could be assessed by examining the area of clearance on gluten containing agar [12]. Furthermore, *ex vivo* culture of the MAM directly with gluten could allow for the assessment of microbial gluten degradation through mass spectrometry analysis. These approaches may provide further insight into the mechanisms underpinning the change in immune responses following gluten ingestion in mice who have been treated with antibiotics.

In conclusion, this study found that altering the microbiome through antibiotic treatment was sufficient to impair the immune response to gluten in mice. The low-grade inflammation observed aligns with the pathophysiology of NCGS. This work suggests that the microbiome may determine the capacity for gluten to induce an immune response and offers a valuable insight into the mechanism underlying NCGS.

## Data Availability

The16S rRNA amplicon and MGS raw sequence reads were deposited under NCBI BioProject accession number PRJNA1234692. The NCBI SRA record is accessible at the following link: http://www.ncbi.nlm.nih.gov/bioproject/1234692.

## Author contributions

JCP designed research, performed experiments, analysed data, interpreted results, prepared figures and drafted the manuscript. ECH supervised research, analysed data, interpreted results, edited and revised manuscript. WSS, SF, SC, KM, SS, CN and HM assisted with experiments. KD, KEH and NJT supervised research, edited and revised manuscript. SK and GLB conceptualised and supervised research, interpreted results, edited and reviewed manuscript.

