## Supplementary data for "Microbiota Modulation Induces Elevated Duodenal Eosinophils Upon Gluten Exposure in Mice: Implications for Non-Coeliac Gluten Sensitivity"

### Slide 1
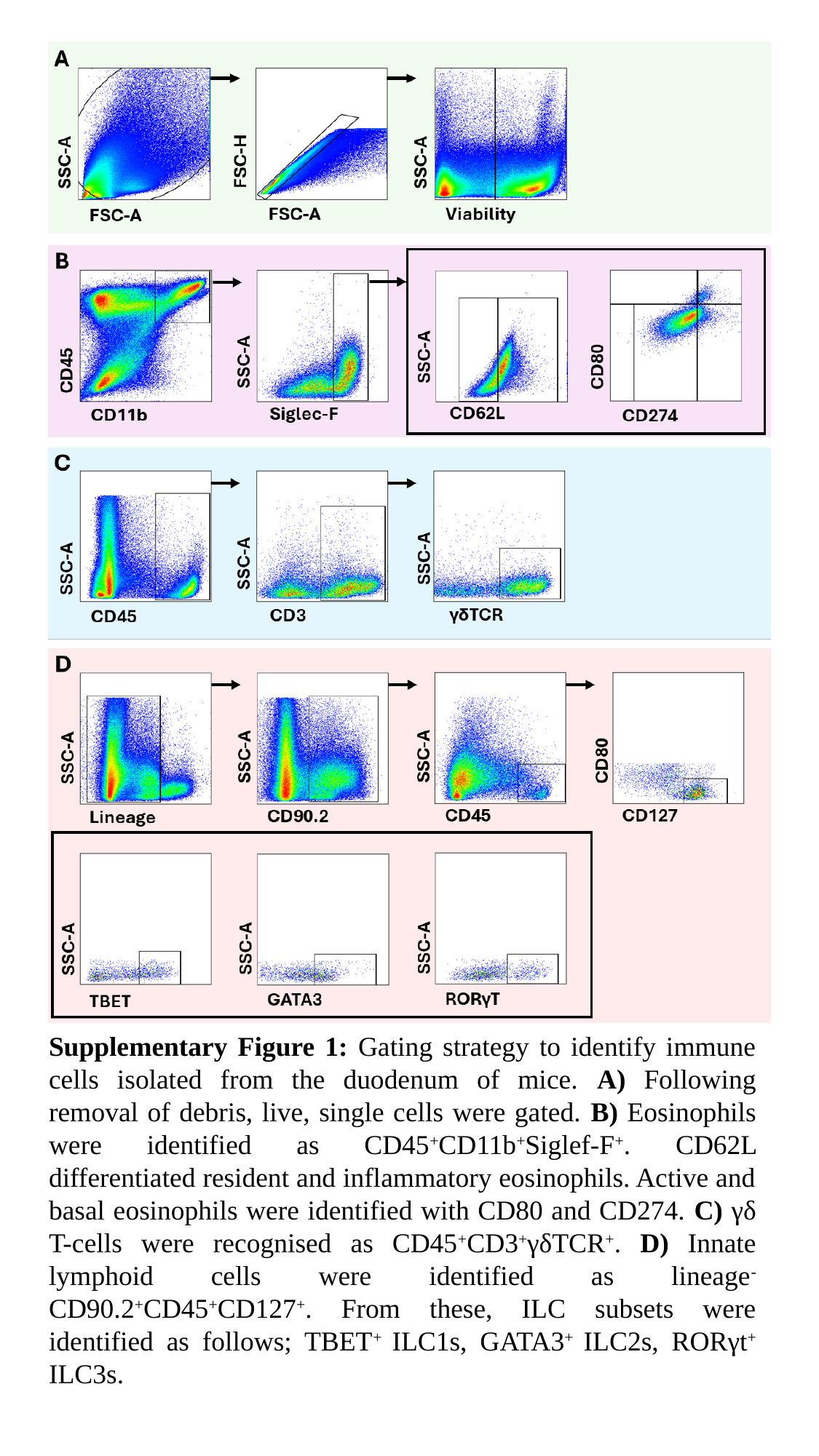

Supplementary Figure 1: Gating strategy to identify immune cells isolated from the duodenum of mice. A) Following removal of debris, live, single cells were gated. B) Eosinophils were identified as CD45^+^CD11b^+^Siglef-F^+^. CD62L differentiated resident and inflammatory eosinophils. Active and basal eosinophils were identified with CD80 and CD274. C) γδ T-cells were recognised as CD45^+^CD3^+^γδTCR^+^. D) Innate lymphoid cells were identified as lineage^-^CD90.2^+^CD45^+^CD127^+^. From these, ILC subsets were identified as follows; TBET^+^ ILC1s, GATA3^+^ ILC2s, RORγt^+^ ILC3s.

### Slide 2
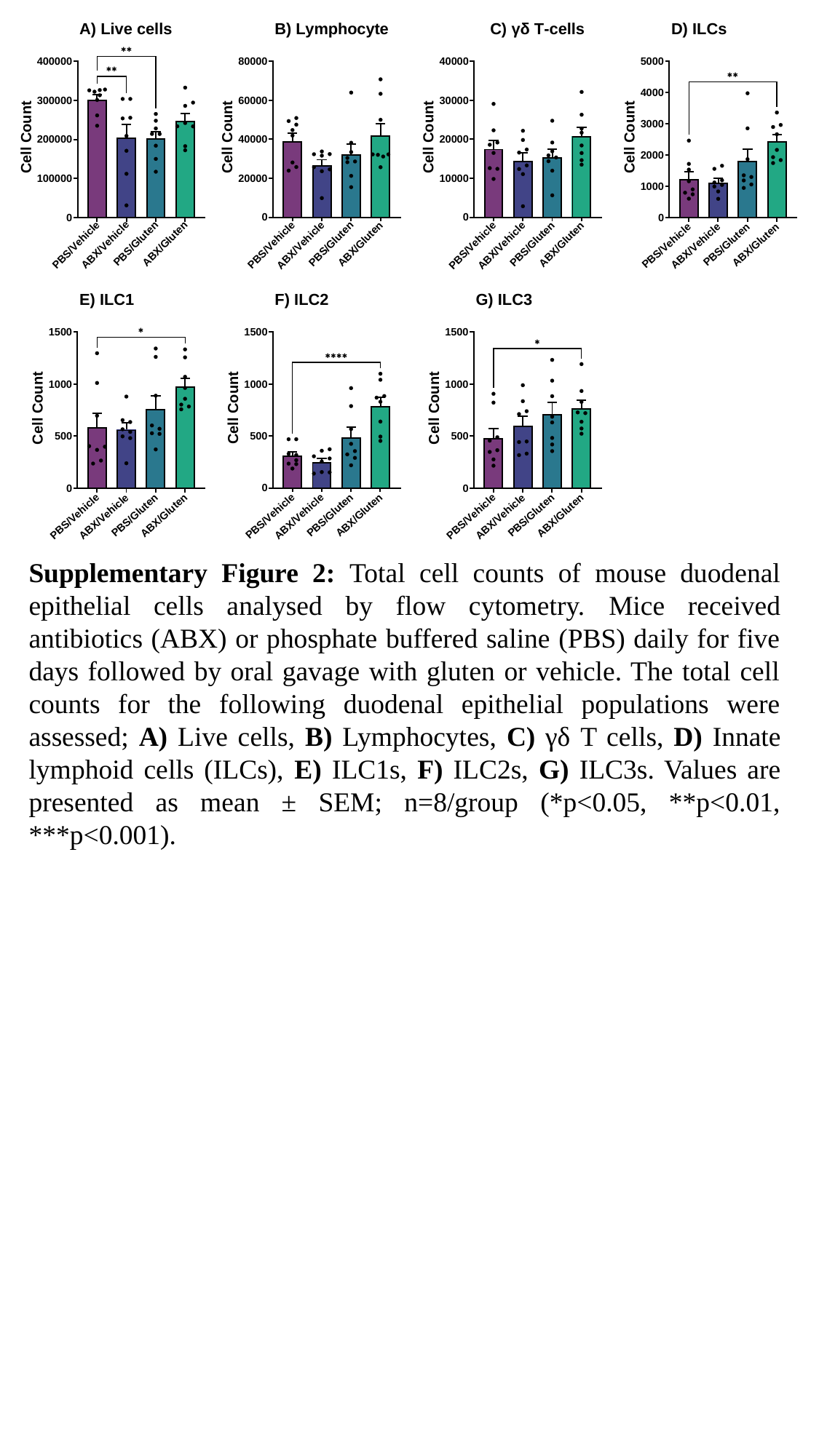

Supplementary Figure 2: Total cell counts of mouse duodenal epithelial cells analysed by flow cytometry. Mice received antibiotics (ABX) or phosphate buffered saline (PBS) daily for five days followed by oral gavage with gluten or vehicle. The total cell counts for the following duodenal epithelial populations were assessed; A) Live cells, B) Lymphocytes, C) γδ T cells, D) Innate lymphoid cells (ILCs), E) ILC1s, F) ILC2s, G) ILC3s. Values are presented as mean ± SEM; n=8/group (*p<0.05, **p<0.01, ***p<0.001).

### Slide 3
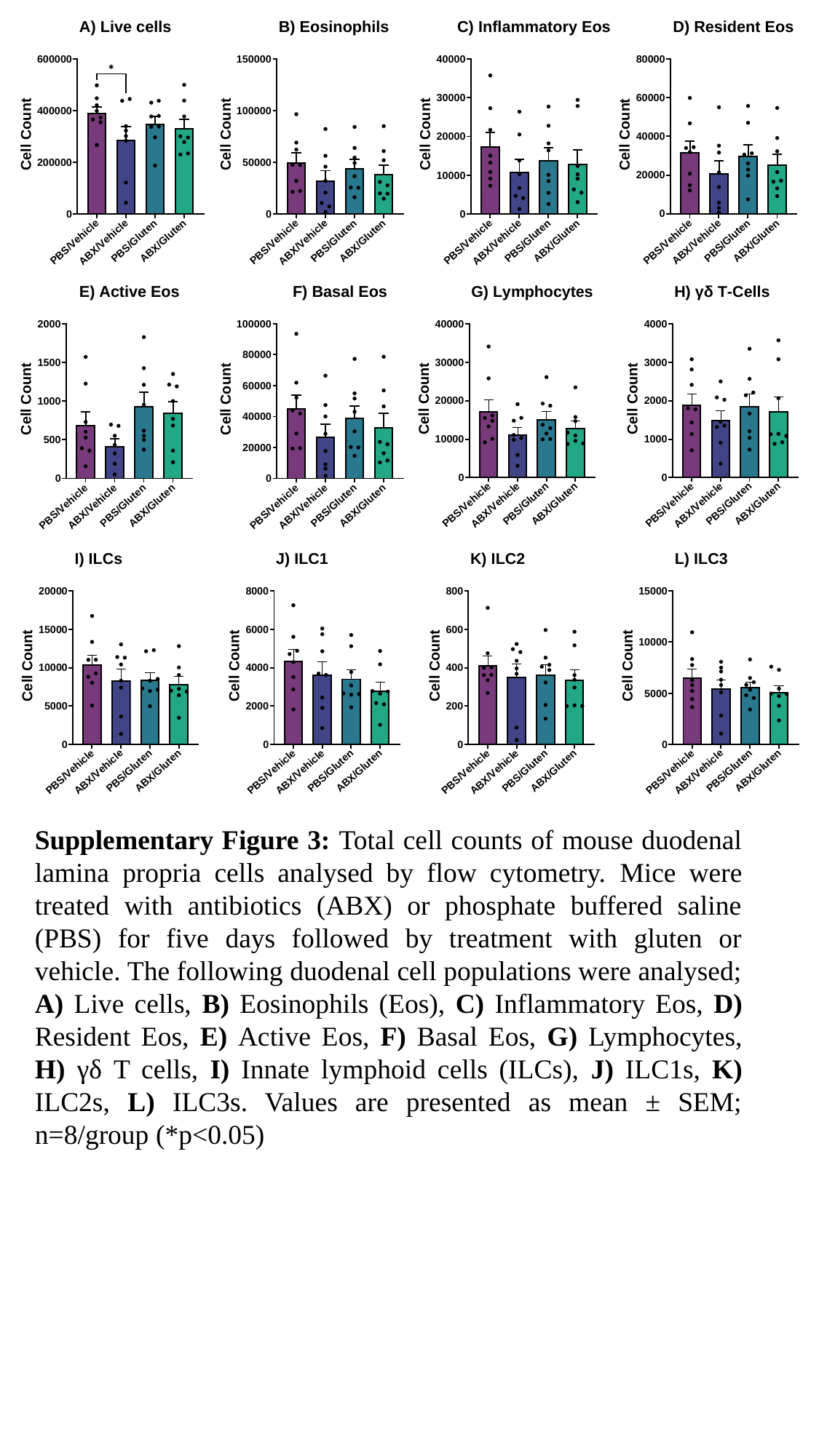

Supplementary Figure 3: Total cell counts of mouse duodenal lamina propria cells analysed by flow cytometry. Mice were treated with antibiotics (ABX) or phosphate buffered saline (PBS) for five days followed by treatment with gluten or vehicle. The following duodenal cell populations were analysed; A) Live cells, B) Eosinophils (Eos), C) Inflammatory Eos, D) Resident Eos, E) Active Eos, F) Basal Eos, G) Lymphocytes, H) γδ T cells, I) Innate lymphoid cells (ILCs), J) ILC1s, K) ILC2s, L) ILC3s. Values are presented as mean ± SEM; n=8/group (*p<0.05)
